# When Online Citizen Science meets Teaching: Storyfication of a science discovery game to teach, learn, contribute to genomic research

**DOI:** 10.1101/2022.05.23.492810

**Authors:** Chrisostomos Drogaris, Alexander Butyaev, Elena Nazarova, Roman Sarrazin-Gendron, Harsh Patel, Akash Singh, Brenden Kadota, Jérôme Waldispühl

## Abstract

In the last decade, video games became a common vehicle for citizen science initiatives in life science, allowing participants to contribute to real scientific data analysis while learning about it. Since 2010, our scientific discovery game (SDG) Phylo enlists participants in comparative genomic data analysis. It is frequently used as a learning tool, but the activities were difficult to aggregate to build a coherent teaching activity. Here, we describe a strategy and series of recipes to facilitate the integration of SDGs in courses and implement this approach in Phylo. We developed new roles and functionalities enabling instructors to create assignments and monitor the progress of students. A story mode progressively introduces comparative genomics concepts, allowing users to learn and contribute to the analysis of real genomic sequences. Preliminary results from a user study suggest this framework may help to boost user motivation and clarify pedagogical objectives.

## Introduction

Biology and evolution inspired the development of many video games, among which we find Spore [1], Plague [2], Cell to Singularity [3] and Niche [4]. The success of these games motivated the development of new approaches for supporting the teaching of biology through video games. Immune Attack [5] and Immune Defense [6], Tyto Online [7], or Genomics Digital Lab [8] have thus been developed to enhance the student experience through practice of key concepts presented in the game. Remarkably, the integration of this technology into classrooms has been shown to help students improve their understanding of biology [9] and performance on tests [10].

The enthusiasm of the public for video games with a scientific narrative also motivated the development of Science Discovery Games (SDG), using real scientific data and designed to engage the participants into research [11]. In the last decade, Foldit [12], Phylo [13], EteRNA [14] and EyeWire [15] gathered hundreds of thousands of online citizen scientists to contribute to the analysis of biological data. Their development offered an opportunity to revisit the visualization and manipulation of scientific data through game design techniques. The novelty and quality of these new interfaces prompted the repurposing of SDGs into professional scientific software [16]. It also very quickly attracted the attention of scientific instructors who saw in these SDGs an opportunity to mobilize students and allow them to engage research while learning [17, 18].

This contribution focuses on our SDG Phylo (http://phylo.cs.mcgill.ca), which launched on November 28^th^, 2010. Phylo aims to engage the public into comparative genomics research through a casual tile matching puzzle game encoding an instance of a multiple sequence alignment problem. The default game board is a 10×25 grid in which we display fragments of genomic sequences as sequences of tiles of 4 different colors representing the 4 nucleotides (i.e., A, C, G, T/U) of a DNA or RNA molecule. Participants move the tiles horizontally but cannot change their order, and their objective is to find a configuration that maximizes a score representing the quality of the alignment of the sequences. At the end of the game, the solutions to these puzzles are stored in our server and later used to help us improve the accuracy of computer-generated alignment of biological sequences stored in public databases (e.g., UCSC genome Browser, Rfam).

The multiple sequence alignment is one of the earliest and still one of the most powerful and widely used computational technique to study the evolution, structure and function of DNA, RNA and protein sequences [19]. In its simplest form, the problem consists in computing a mapping of nucleotides (or amino acids) between several biomolecules that respects the order in these sequences. Yet, the computation of a reliable alignment faces many challenges. Given a scoring function, aligning multiple sequences is a computationally hard problem [20]. Worse, the definition of a reliable objective function to estimate the quality of an alignment remains an unsolved challenge [21–24]. Despite major innovations and undeniable progresses across the years, the problem is still open [25]. Fully automated alignments often contain inaccuracies that require an expert to manually curate and fine-tune the alignment. Noticeably, popular alignment databases such as Rfam [26] are already semi-automatically collecting improved alignments submitted by their users [27].

The game is fast, simple, and can be played by any user with no prior knowledge of biology (See Figure 1). It does not require registration to participate, which lowers the barrier of entry and increase participation making it an effective tool for teaching activities.

**Fig. 1.**
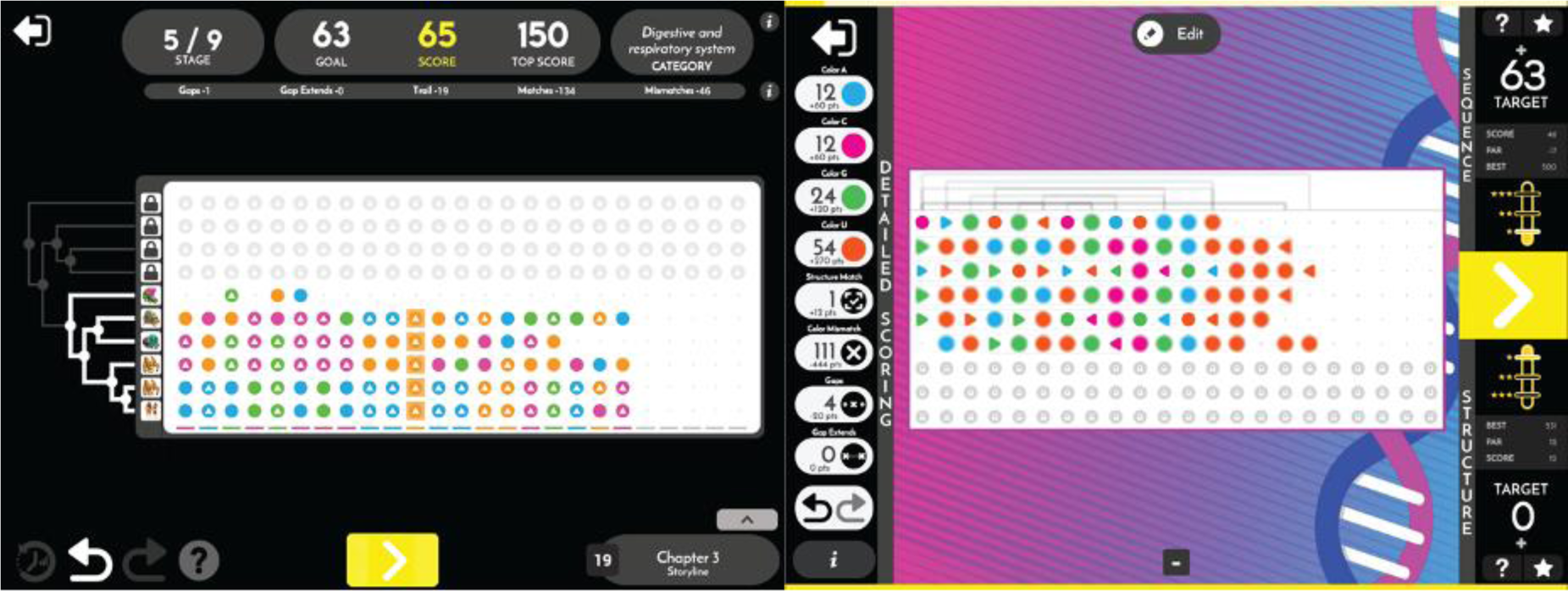
User interfaces of the scientific discovery games. (left) Phylo DNA puzzle interface featuring a phylogenetic tree. (right) Ribo RNA puzzle interface with base pairs represented as arcs on the top of the game board.

Since 2010, hundreds of thousands of users have played Phylo. It seemed to us unreasonable to concentrate all this public participation into our sole hands. Therefore, in 2013, we opened up our platform, allowing external researchers to use our system to solve their sequence alignment problems through crowdsourcing with OpenPhylo [28]. To our knowledge, it is the first time that access and full control of an SDG has been shared with the entire scientific community. OpenPhylo enables scientists to upload their own sequences which are converted to puzzles and introduced to gamers through the Phylo game. Scientists can then track the number of times their puzzles have been played and download at any time an alignment of the sequences improved with the solutions collected by Phylo.

The objective of this work is to bring OpenPhylo to the next level, where our aim is to support scientists, instructors and power users of Phylo. Instructors can now use the OpenPhylo platform to create Phylo assignments for students to allow them to learn and contribute to real scientific data collection. Phylo power users can use the OpenPhylo website to view advanced statistics completed on their account throughout their playtime in Phylo. But most importantly, we fully redesigned the gameplay and user experience to fulfil the needs of teaching applications. We introduce a story mode that builds a narrative connecting the scientific activities (i.e., the puzzles) together. The story enables us to progressively present key scientific concepts and tutorial puzzles allowing the participants to practice basic techniques on synthetic data before applying them on real scientific puzzles. The story is also punctuated with optional quizzes that diversify activities and could be used to test scientific knowledge of participants. Simultaneously, the inclusion of educational content also aims to increase engagement of participants [29].

This contribution describes a generic strategy and series of recipes to streamline the use of SDGs as teaching support tools, and therefore expand the integration of citizen science activities in classrooms [30]. In practice, we implemented our strategy in our genomic SDG Phylo. We conducted a small user study in a bioinformatics class to validate the relevance and effectiveness of our approach. Our results suggest that the storyfication of SDGs is a promising approach to deploy citizen science activities in class and promote the engagement of students. We also find that it can help to clarify the presentation pedagogical objectives.

## Methods

OpenPhylo was designed to be a meeting point between citizen scientists (i.e., a player of the SDG), scientific instructors, and scientific experts (See Figure 2). All three user types can log in to our system and access the services on the web portal located at http://openphylo.cs.mcgill.ca. In this section, we describe the new teaching-related features, as well as general improvements made to Phylo that benefit all users.

**Fig. 2.**
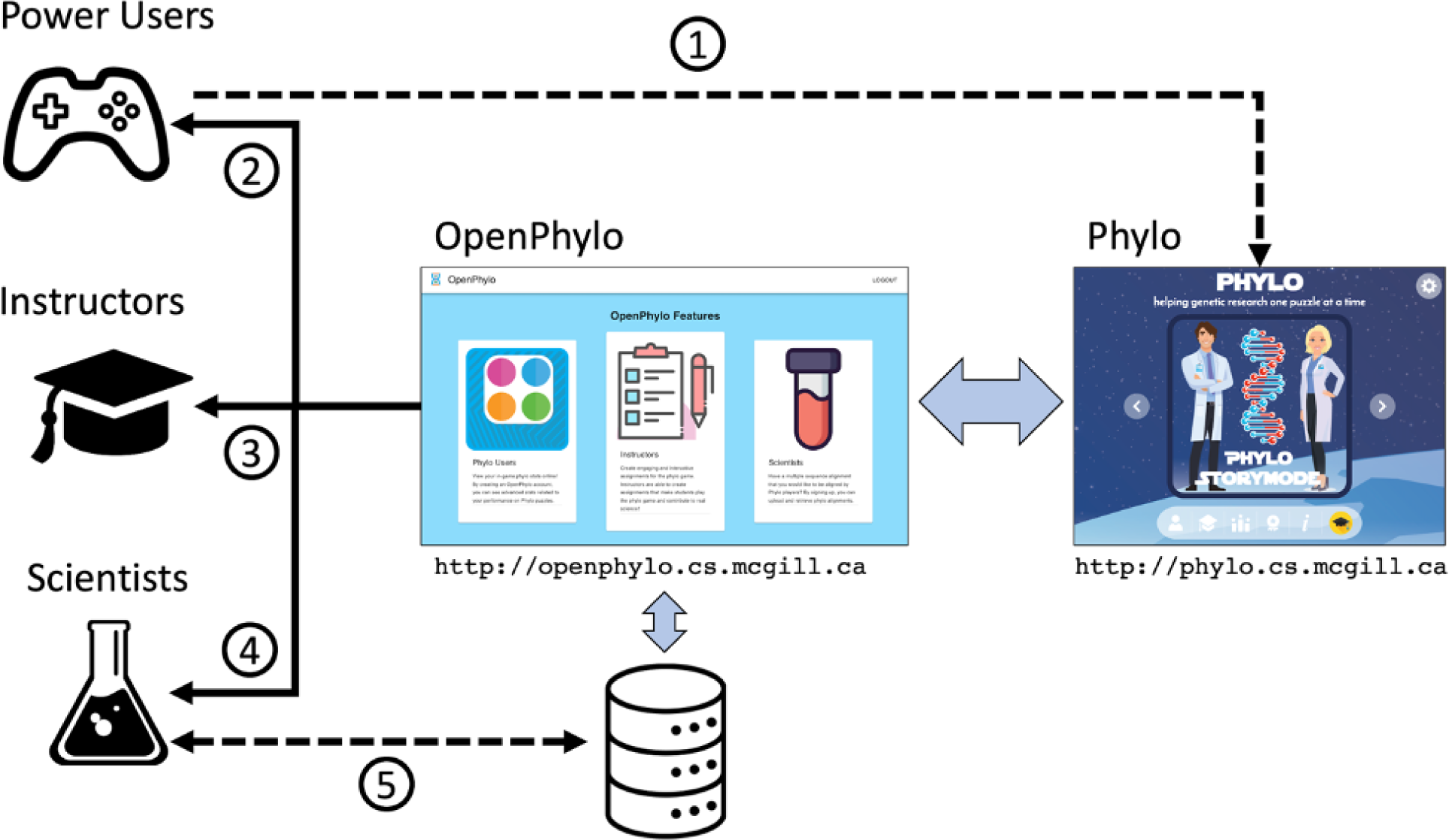
Phylo and OpenPhylo system overview. (1) Power users (a.k.a. citizen scientists) play Phylo, and (2) connect to OpenPhylo to access personal statistics. (3) Instructors connect to OpenPhylo to create assignments and invite power users. (4) Scientists connect to OpenPhylo to (5) upload sequences to the puzzle database and download solutions from participants.

### New teaching features Story mode

Phylo puzzles are now integrated into a story and a roadmap broken down into chapters (See Figure 3). In previous versions of Phylo, participants requested puzzles based on a difficulty level or associated disease. Each puzzle was independently played from the others. The new version of Phylo lets participants engage with the SDG through a story introducing puzzles by increasing difficulty levels. The story serves several purposes, including making the activity more fun and entertaining, but also enabling us to progressively introduce key concepts for computing accurate sequence alignments such as the scoring of mutations and insertions-deletions (indels), the use of phylogenetic trees, or covariation in RNA alignments. It also allows to seamlessly walk the players through a tutorial describing techniques and giving tips for building good alignments, and it makes it easier to monitor progress. Finally, it’s a flexible framework to introduce other education activities such as quizzes.

**Fig. 3.**
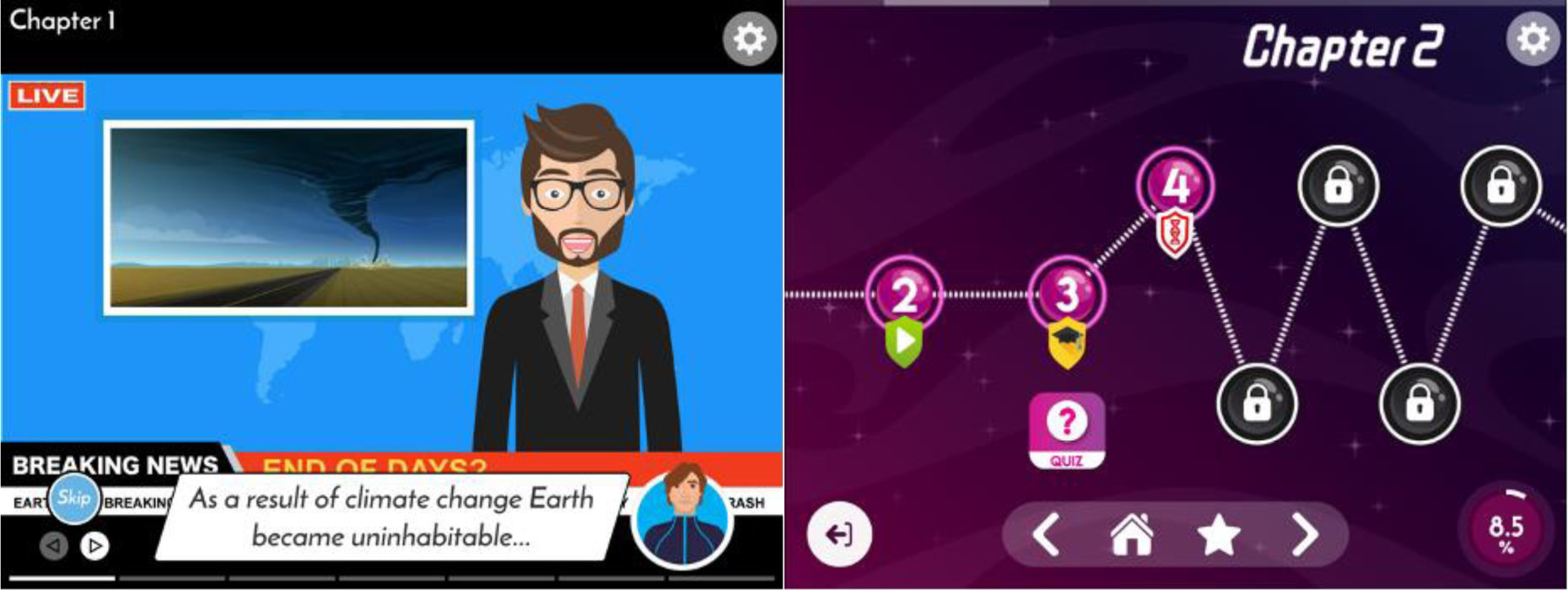
Story mode. (left) a slide telling the story in which we embed the Phylo and Ribo puzzles. (right) roadmap of the story line. Each node along the path represents either a series of slides telling part of the story, a tutorial, a quiz, or a puzzle. The bottom menu allows the player to navigate the roadmap. A progress wheel can be found on the bottom right of the screen.

The story is based on a science-fiction scenario where humans try to re-colonize earth after leaving it a long time ago because of environmental issues. This plot aims to bring more purpose to the activities but also to highlight the complexity and broad impact of environmental challenges. Yet, the humor and far future timeline intends to prevent dramatization that could impact the pedagogical objective of our SDG.

This story mode was the first step required to turn Phylo from a casual game to an educational process with clear objectives and a sense of progress observable by both the user and the teacher.

### Quizzes

To complement the learning process embedded in the story, a new types of activity, quizzes, was added to Phylo (see Figure 4). Unlike the puzzles, the answers to the quizzes submitted by the participants are not used to annotate or classify any scientific data. However, they offer an additional tool for instructors who are using Phylo as a teaching support, and they increase the diversity of the tasks, which enriches the user experience and overall entertainment value of the SGD.

**Fig. 4.**
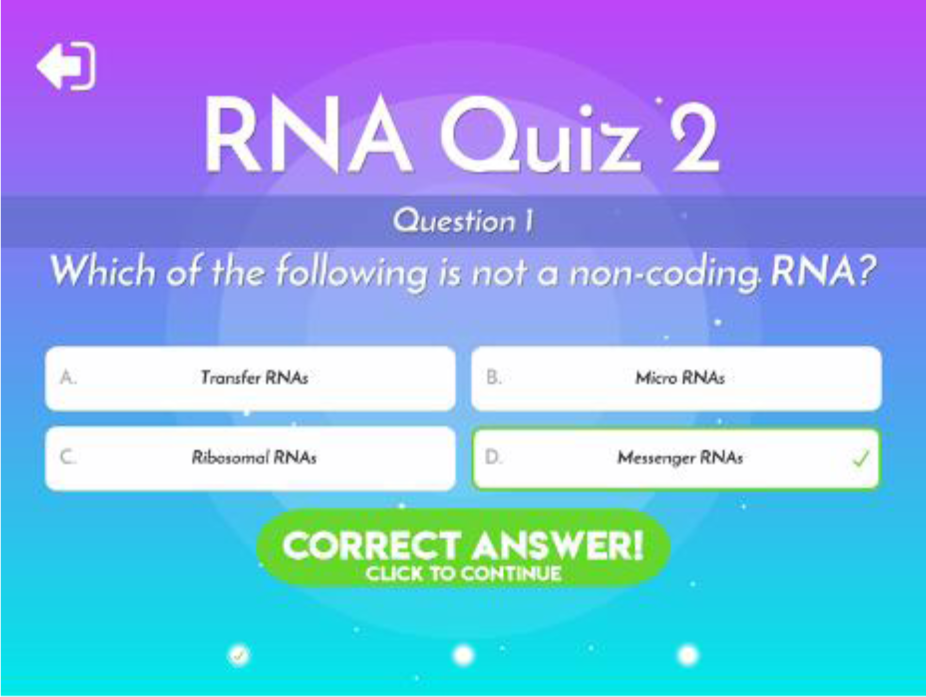
Example of an optional quiz presented to the participants before starting a series of puzzles.

Currently, quizzes are optional and offered to the player before starting the citizen science activities (i.e., the puzzles) as they advanced through the story mode. The questions are related to the current chapter of the story, but not necessarily directly connected. They aim to test general knowledge in the field of genomics rather than specific concepts introduced in the game. Nevertheless, they can be customized by the instructor. A full description of the questions available in Phylo is available in the supplementary material.

### Courses and Assignments

Along with the story mode and the quizzes, courses and assignment modules have been added to Phylo. A course is a general structure in which instructors can register their students and organize several assignments on a particular topic. Assignments are sets of Phylo-related tasks for students that can be monitored by instructors.

In OpenPhylo, there are two types of assignments: story mode and classic mode.

The story mode assignments leverage the new story mode and quizzes by asking students to complete specific chapters of the story, which make the students go through hand-picked puzzles and quizzes to guide them through an appropriate difficulty scaling.

Classic assignments revolve around Phylo’s classic mode where users select a disease type and solve random puzzles.

The instructors can define the difficulty level and the number of puzzles they expect students to complete. They allow students to explore and play the Phylo game without constraints.

### A new user type: instructors

At the time of creating an account, all users are considered power users (i.e., regular players). After creating an account, they can upgrade to a scientist role, or the brand-new instructor role.

Instructors have several unique functionalities. The first one is the ability to create courses through a dedicated form. Each course has a name, a description and requires a list of student emails. Once an instructor has created a course, they can then access that course’s page. On each course page, instructors have the option to edit a course’s details, add assignments and see a list of past, ongoing, and upcoming assignments. The edit course page allows instructors to change the name and description of a course and also add or remove students. Students are regular Phylo users enrolled into a course by the instructor.

An instructor can create assignments through a button on the home page of a course. They are then presented with a form that asks for an assignment name, description, start date, end date and assignment type (i.e., classic or story-based). Once an assignment is created, OpenPhylo sends an email to each student in the course on the start date of this assignment (See Figure 5). Then, each student can accept this request and start playing Phylo to complete this assignment. A student already owning a Phylo account can simply use the same account if the email address corresponds. Students without pre-existing Phylo accounts must register with their student email. Once an assignment is started, instructors can monitor in real-time the status of each student. Figure 5 shows a status page for a story mode assignment. Instructors can check whether students have accepted the assignment and which tasks they have completed.

**Fig. 5.**
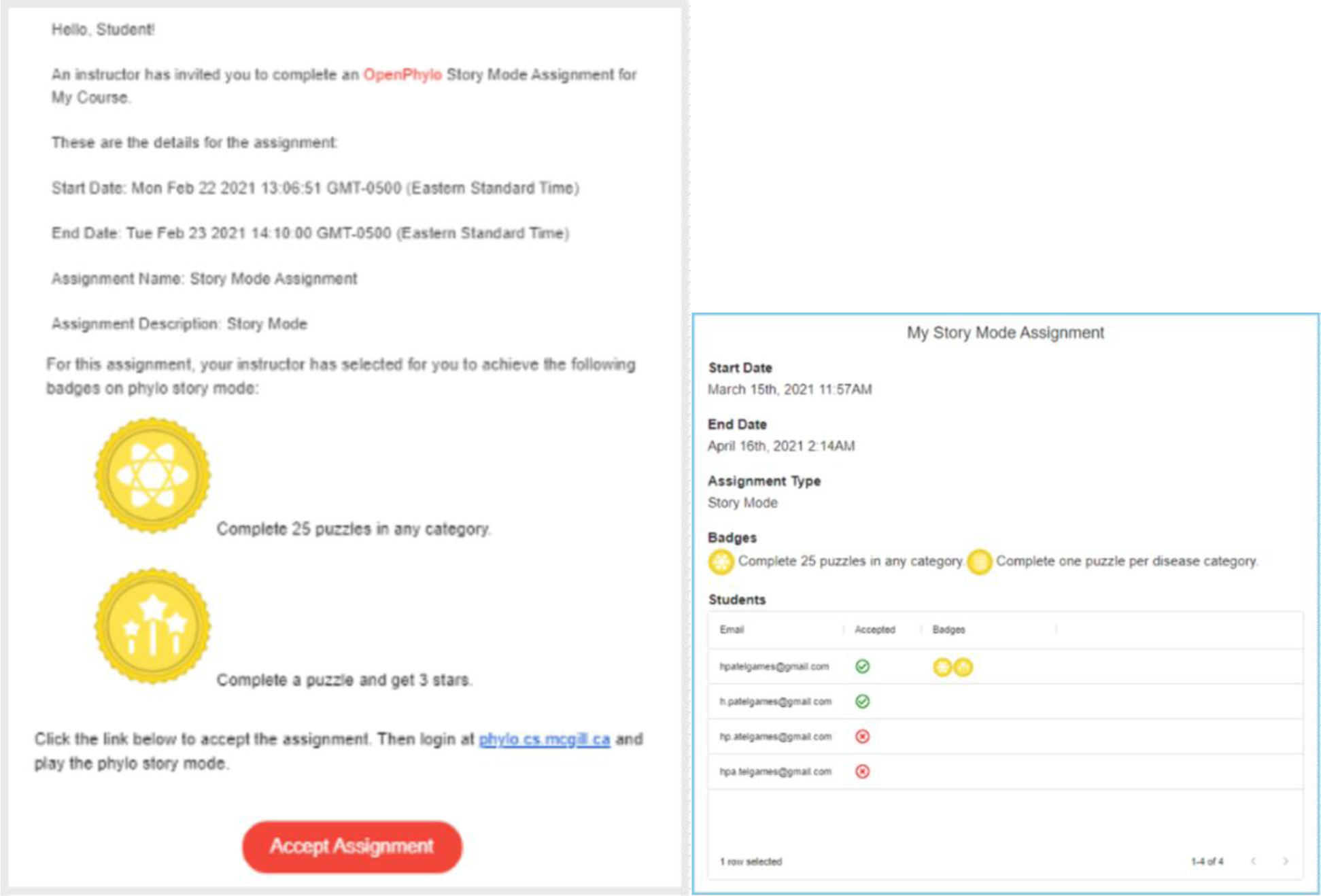
Example of an assignment. (left) Automatic message with instructions send to the students invited by the instructor. (right) information panel allowing the instructor to monitor the progresses of the participants to the assignment.

To accommodate the use of Phylo and OpenPhylo to different contexts beyond the pre-programmed questions covering genomics and general bioinformatics, we allow an instructor to customize story mode assignments, which include the possibility of building a specific level map and specific quizzes.

Thus, instructors have all the tools they need to adapt Phylo to their specific teaching needs.

### Scientist users

The scientist user role was first introduced with the initial release of OpenPhylo in 2013 [28], but has been improved in this new version to lower the barrier to entry.

Scientist users can upload their multiple sequence alignments into Phylo, and then, once they are played by enough users, receive alignments that can improve on existing state of the art alignments done with tools such as MUSCLE [31].

Like instructor accounts, scientists are a special user type requiring an additional registration form to apply to be a scientist, which is sent to the administrators of OpenPhylo to validate the new role. This validation mechanism reduces the risks of a malicious user flooding the system with corrupted data.

Figure 2 showcases the workflow for a scientist to create puzzles and collect the solutions. The first step for a scientist is to upload an unaligned FASTA file containing the sequences that they wish to align. Each file must include a maximum of 10 sequences. In previous work, we developed aggregation methods to align more than 10 sequences, but we decided to not implement those in OpenPhylo because of the experimental nature of this work and the diversity of the parameter space; we wanted to prioritize the quality of results for now. Once the FASTA file is uploaded to OpenPhylo, the backend runs a python script that aligns the input sequences with MUSCLE. [31]

Some Phylo variants require a phylogenetic tree. In the past scientists were required to upload a tree with their sequences. Now, it is optional. In absence of a tree, OpenPhylo uses IQ-Tree to compute a phylogenetic tree. [32]

Once the initial alignment is created, the puzzles can be generated. First, the scientist specifies the number of puzzles to create. We use heuristics developed in previous studies to determine which regions of the full alignment could be improved with Phylo [28]. Since the maximum length of a sequence in a basic Phylo puzzle is 15, we scan our full alignment with a 15 nucleotide-long window and extract the most promising regions. Currently, OpenPhylo uses a simple heuristic selecting regions with the largest number of gaps and mismatches, which suggest there is room to improvement the alignment.

Once a region is selected, a puzzle is automatically created. Then, a recursive function is called on the left and right side of the region selected to create additional puzzles. This allows us to increase the coverage of this region while preventing overlaps. The process stops when we reach the maximum number of puzzles requested by the user.

After creation of the puzzles, the scientists can monitor the statistics for the puzzles they created (e.g., number of puzzles created, coverage of the alignment, alignment scores). OpenPhylo calculates sequence specific metrics such as the number of puzzles created, total sequence length, and puzzle coverage percentage. These numbers allow the scientist to get an overview of how much of their sequence is covered by Phylo puzzles. The player-improved alignment is obtained by aggregating the best solution to each puzzle created and replacing the region in the original alignment with the solution. OpenPhylo also computes an alignment score for both the original and player-improved alignments, using MUSCLE’s sum-of-pairs scoring.

### General improvements to Phylo

#### Puzzles and scoring

We developed expanded the diversity of sequence alignments task to progressively expose the users to various concepts and facilitate the learning process.

This new release integrates our previously released SDG Ribo (See Figure 1), which asks the user to align structured RNAs (i.e., with a secondary structure annotation) [33] rather than pure sequences. In the same spirit, we developed simplified and advanced versions of Phylo and Ribo that allow users to practice separately different scoring mechanisms and features of the games.

Indeed, we now distribute two versions of Phylo. The first one is a plain multiple sequence alignment scored with an affine gap model. Match (+1), mismatch (-1) and indel costs (-2 of a gap opening and -1 for a gap extension) are identical to those used in previous versions of Phylo [13, 28], where parameters are rounded for transparency and better interpretability by the users. [34] In contrast to the original SDG, this version does not use a phylogenetic tree but a sum of all pairs scoring model, which aims to lower the barrier of entry for new users. The other version of Phylo distributed on our platform is the classic one that uses a phylogenetic tree and a customized version of the Fitch algorithm for the solving minimum parsimony problem [35].

The game also features two versions of Ribo. The first one aims to align RNA sequences with secondary structure annotations. The score is the sum of a sequence score computed as a sum-of-pairs and a structure score rewarding each aligned base pairs. In the other version, the scoring scheme is identical, but the user can edit the base pairing annotation. These variants of the multiple sequence alignment problem enable us to progressively introduce users to complex scoring schemes, allowing them to practice on simpler problems before moving on to more sophisticated ones.

Over the course of the story, these different types of puzzles (see General improvements) are introduced to the player in order of increasing complexity. The chapters that introduce a new type of puzzle (i.e., chapters 2, 3, 6, and 7) include a short tutorial and a couple of synthetic practice puzzles before solving puzzles using real data. Furthermore, we developed a hint mechanism that suggests move to players needing assistance, which also aims to improve engagement.

The performance of participants per puzzle is rated using a three-star scoring system which provides additional motivation to complete tasks at a higher level of accuracy and reinforce the acquisition of the concepts introduced in the SDG [36]. The number of stars attributed to the players is determined by its similarity to the highest score obtained so far. A higher-than-ever score receives three stars, whereas a score equal to the baseline (i.e., the “Par” score that is obtained as the score of the alignment computed by automated methods) receives one star.

#### Badges

Badges are collectible items symbolizing milestones. Participants that successfully complete activities in a chapter featuring puzzles receive badges associated with the chapter’s theme. Other badges are delivered for global achievements such as total puzzles successfully completed or first time a user obtains a 3-star rating. To avoid redundancy, we do not enable instructors to create their own new badges. However, we accept requests to create new ones.

Properly implemented, the representation of achievements is a powerful source of motivation for participants [37] that can boost user retention and motivation in learning games. For an educator, it also is a convenient tool to define the objectives and track the progress of the students. A list of the badges currently available in Phylo is available in the supplementary material.

## Results

We conducted a user study from March 27th, 2022, to April 26th, 2022, with 13 students registered in a graduate level bioinformatics course. The 13 students, 11 of which were undergraduate students, were asked to complete Phylo’s story mode. We created an assignment using the instructor functionalities, tasking them with completing the entire story mode. They all completed the entire story during the time allocated and collected all badges associated with chapters 2, 3, 6, 7, and 8 (see supplementary material).

Then, we asked the students to complete an anonymous survey to describe their experience and rate various functionalities of the game. The participation in the survey was optional and was not affecting the grading in the course in any way. Yet, all students accepted the invitation and provided anonymous feedback.

First, we asked about the benefits of implementing a story mode. We were interested in assessing whether the story is entertaining (Q1), instructive (Q2), helps to connect activities together (Q3), and offers an additional incentive to complete the tasks (Q4). Figure 6 shows our results. On all aspects, the feedback was globally positive. The strongest agreement was obtained on the first question “The activity is fun”, where 6 participants strongly agreed, 4 agreed, 1 was neutral and 2 disagreed. It validates our primary objective that is to make citizen science and teaching activities more entertaining. Though, the 2 disagreements suggest the approach does not suit all participants. On the fourth question “the story gives more motivation”, 4 participants strongly agreed, 7 agreed, and 2 were neutral. Notably, this is the only question where no one disagreed. This result suggests that the story improves engagement. A similar level of agreement was obtained with the third question “the story It helps bridging puzzles together”, where 4 participants strongly agreed, 7 agreed, 1 was neutral and 1 disagreed. Answers to the second question “The story is instructive” are also positive albeit slightly more mitigated. 4 participants strongly agreed, 5 agreed, 2 were neutral and 2 disagreed. It confirms the story effectively introduces key concepts.

**Fig. 6.**
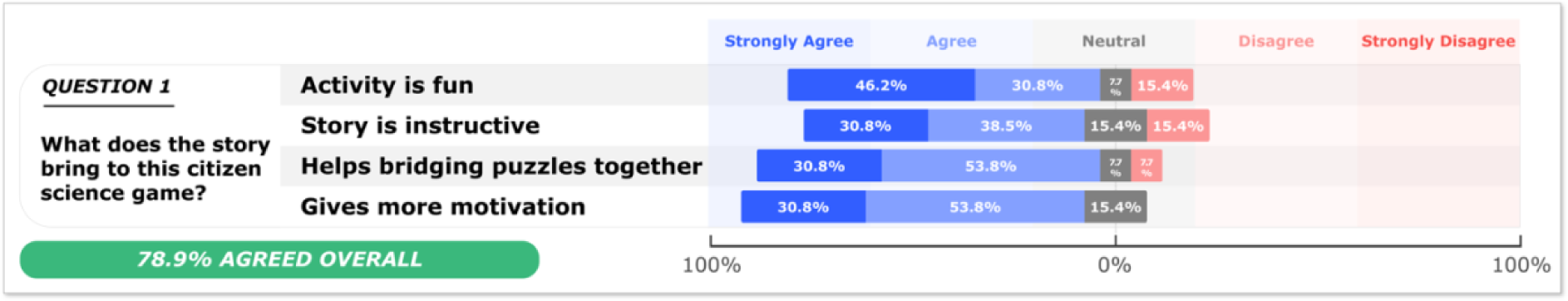
Results of the user survey about the impact of the storyfication of Phylo.

For each chapter that introduced a new type of puzzle, we aimed to assess the relevance and educational impact of this presentation of the citizen science activities. We asked the participants if the “tutorial was helpful” (Q1), “the practice puzzles help to understand the task” (Q2), “the puzzles are fun” (Q3), “the 3 stars rating motivates the player to get better alignment” (Q4), and if “the pedagogical objective is clear” (Q5). The results are presented in Figure 7. The answers are consistent among all chapters, and noticeably consistent for DNA (Phylo) and RNA (Ribo) chapters.

**Fig. 7.**
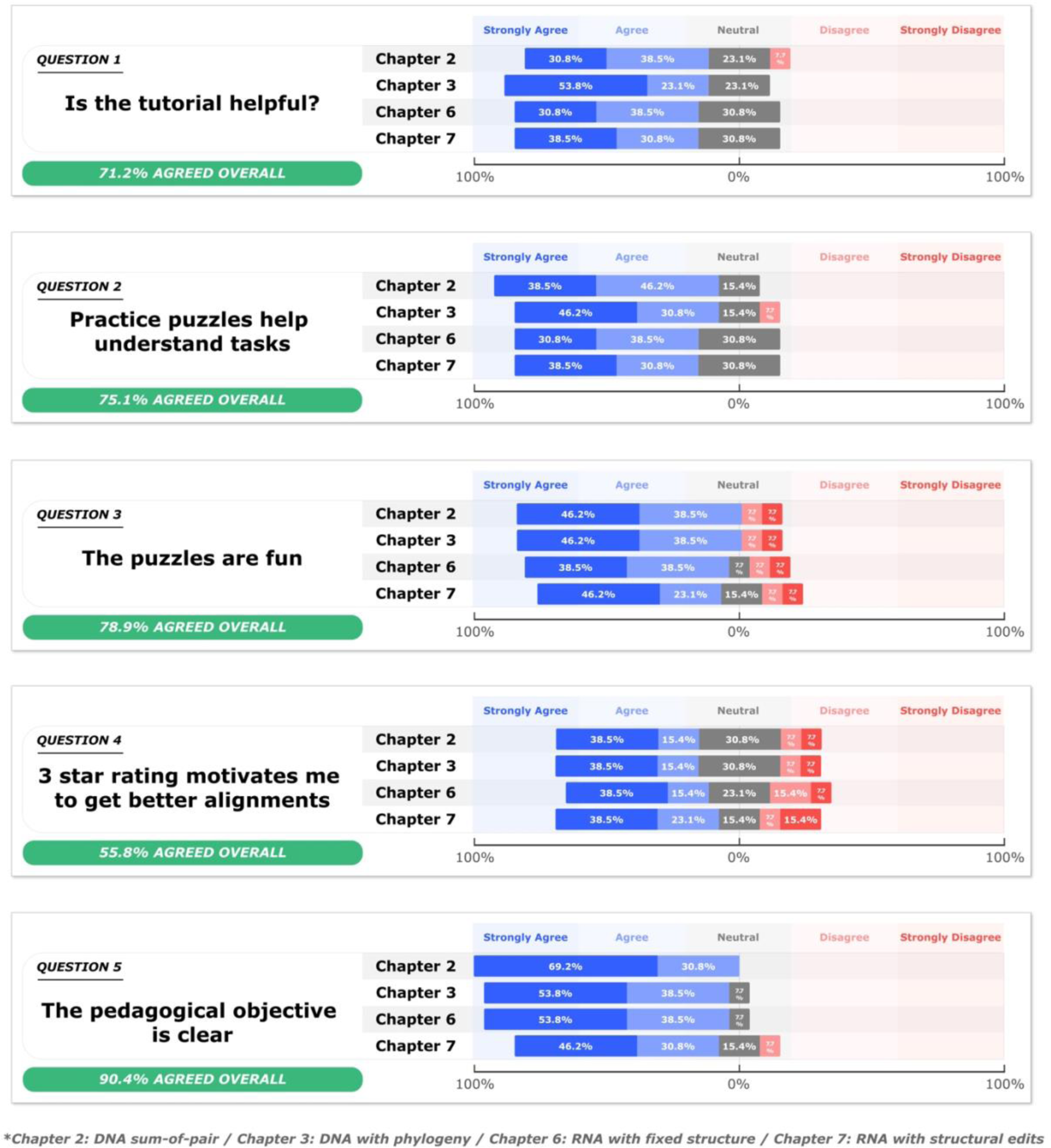
Results of the user survey about the educational impact of each chapter.

The strongest agreement was obtained on the fifth question (i.e., clarity of the pedagogical objectives), which supports the effectiveness of our approach. We note a slight decrease in the agreement in the last chapter, but the drop is modest and correlates with the complexity of the tasks.

Strong agreements were also observed on questions 1 and 2. These two questions aimed to understand how useful are the tutorial and practice puzzles that have been introduced in this new version of Phylo. Our results suggest that both are equally valuable features. Answers to the third question are also very positive but exhibit more variance. This is in agreement with previous observations and confirms that the activity is fun for the majority of participants but does not necessarily suit everyone. Only question 4 inquiring about the usefulness of the 3 stars rating system was more mitigated. Though, previous citizen science games already validated the effectiveness of this system [36], which led us to think that the benefits of this scoring mechanism could be improved in the story mode.

Finally, it is worth noting that some optional comments mentioned the background music, including one participant who reported that it helped to play more puzzles. This observation suggests that music could be leveraged to promote engagement. [38]

## Discussion

OpenPhylo aims to become a crossroad for citizen scientists, educators, and genomics researchers. We designed a simple web interface to enable users to complete basic requests including accessing statistics, create assignments and invite participants, or use Phylo to ask the citizen scientists to improve an alignment. To achieve this objective, we also embedded the SDGs Phylo and Ribo in a story game where participants can progressively learn the principles of sequence alignment and key concepts in comparative genomics and evolutionary biology. This new release is a major step in the evolution of our SDG Phylo. It proposes an integrated view of citizen science in research and education in the internet era. Yet, the platform is still under active development and this work constitutes a proof-of-concept that will be refined in upcoming years. Importantly, we emphasize that the story and instructor tools are not intended to be a substitute but rather an additional tool to support educational activities and engage more participants in genomics.

The results of our user study validate the relevance and effectiveness of our approach. In particular, the story appears to be a powerful vehicle to introduce and explain key concepts in comparative genomics. But this user study also revealed areas of improvement and possible limitations. Among them, we note that a gamified approach might not suit a minority of students more inclined toward classical methods. This observation suggests that participation to citizen science activities in a class should remain optional. However, the fact that the majority of users seem to find the new teaching features enjoyable is very encouraging and appear to suggest that Phylo does not need to be used as a mandatory activity to spark the interest of players.

The three stars rating system also provided less motivation to achieve high scores than we hoped. Although, we believe this aspect could be improved if we make the star ratings more visible and integrated within the story mode. Interestingly, users mentioned features that have a greater impact than we anticipated. In particular, the background music appears to improve engagement. Yet, we acknowledge that it may not be a universal rule and that users should be allowed to customize it. It remains that many other advanced features could be integrated to promote user engagement. [39]

The introduction of a story mode together with the instructor interface created a promising synergy. Our experience suggests that further development should tap into the potential of this system. Currently, Phylo proposes a single and immutable story. The same observation holds true for the quizzes and badges that are predefined. We anticipate that instructors may benefit of customizing or even editing the story, quizzes, and badges for their teaching activities. The development of a mobile version of Phylo is another a major milestone in facilitating access to our SDG and lowering the barrier of entry. This evolution of Phylo is an attempt to be more inclusive and to offer a more equitable access to students who may not have immediate access to a computer.

The scientist interface could also be improved to allow researchers to better exploit the potential of Phylo for improving alignments. This includes the expansion of processing tools (i.e., preliminary alignment of the sequences, different methods of aggregation of the solutions), increase in the number and length of sequences that can be submitted, and more control to create the puzzles. But the most significant improvement could be to automatically grant privileges of scientists to instructors to allow them to create customized puzzles for their students.

Finally, the concepts introduced with OpenPhylo could eventually be expanded to other massive citizen science initiatives such as Borderlands Science [40] to support the expansion of science literacy in our society. [41] We also anticipate that a better integration and more systematic use of SDGs in education could contribute to address challenges in the diversity of participation in citizen science. [42]

## Availability

Phylo is freely available at http://phylo.cs.mcgill.ca or as a mobile application in the Apple store and Google Play. OpenPhylo is also freely available at http://openphylo.cs.mcgill.ca. Scientist users who wish to use OpenPhylo to upload sequences, are manually verified by OpenPhylo administrators.

## Acknowledgements

The authors would like to thank Hilda Olivier and Tim Liu for their contribution to the translation of the game in, respectively, Spanish and Chinese. The authors are also grateful for the feedback, comments, and suggestions from all users of the beta version of Phylo and OpenPhylo and would like to thank all participants to this citizen science initiative. This work was supported by research grants from Genome Canada and Genome Québec.

## Notes

http://phylo.cs.mcgill.ca

### Competing Interest Statement

The authors have declared no competing interest.

### Summary of Updates

Missing middle author in the author list. Minor text changes.

## References

1. Maxis. Spore. How will you create the universe?. 2008; Available from: www.spore.com.

2. Creations, N., Plague Inc. 2012.

3. Lunch, C. Cell to Singularity - Evolution Never Ends. 2018; Available from: www.celltosingularity.com.

4. Studio, S.F., Niche - a genetics survival game. 2017.

5. Scientists, T.F.o.A. Immune Attack. 2008; Available from: www.molecularjig.com.

6. Games, M.J. Immune Defense. 2014; Available from: www.molecularjig.com.

7. Games, I. Tyto online. where students actually do science. 2020; Available from: www.tytoonline.com.

8. Spongelab, Genomic Digital Lab. 2007.

9. Miller, L., et al., An online, interactive approach to teaching neuroscience to adolescents. CBE Life Sci Educ, 2006. 5(2): p. 137–43.

10. Stegman, M., Immune attack players perform better on a test of cellular immunology and self confidence than their classmates who play a control video game. Faraday Discuss, 2014. 169: p. 403–23.

11. Das, R., et al., Scientific Discovery Games for Biomedical Research. Annu Rev Biomed Data Sci, 2019. 2(1): p. 253–279.

12. Cooper, S., et al., Predicting protein structures with a multiplayer online game. Nature, 2010. 466(7307): p. 756–60.

13. Kawrykow, A., et al., Phylo: a citizen science approach for improving multiple sequence alignment. PLoS One, 2012. 7(3): p. e31362.

14. Lee, J., et al., RNA design rules from a massive open laboratory. Proc Natl Acad Sci U S A, 2014. 111(6): p. 2122–7.

15. Kim, J.S., et al., Space-time wiring specificity supports direction selectivity in the retina. Nature, 2014. 509(7500): p. 331–336.

16. Cooper, S., et al., Repurposing Citizen Science Games as Software Tools for Professional Scientists. FDG, 2018. 2018.

17. Dsilva, L., et al., Creating custom Foldit puzzles for teaching biochemistry. Biochem Mol Biol Educ, 2019. 47(2): p. 133–139.

18. Miller, J.A., et al., Introducing Foldit Education Mode. Nat Struct Mol Biol, 2020. 27(9): p. 769–770.

19. Sankoff, D., C. Morel, and R.J. Cedergren, Evolution of 5S RNA and the non-randomness of base replacement. Nat New Biol, 1973. 245(147): p. 232–4.

20. Wang, L. and T. Jiang, On the complexity of multiple sequence alignment. J Comput Biol, 1994. 1(4): p. 337–48.

21. Notredame, C., L. Holm, and D.G. Higgins, COFFEE: an objective function for multiple sequence alignments. Bioinformatics, 1998. 14(5): p. 407–22.

22. Thompson, J.D., et al., Towards a reliable objective function for multiple sequence alignments. J Mol Biol, 2001. 314(4): p. 937–51.

23. Do, C.B., et al., ProbCons: Probabilistic consistency-based multiple sequence alignment. Genome Res, 2005. 15(2): p. 330–40.

24. Loytynoja, A. and N. Goldman, Phylogeny-aware gap placement prevents errors in sequence alignment and evolutionary analysis. Science, 2008. 320(5883): p. 1632–5.

25. Chatzou, M., et al., Multiple sequence alignment modeling: methods and applications. Brief Bioinform, 2016. 17(6): p. 1009–1023.

26. Kalvari, I., et al., Rfam 14: expanded coverage of metagenomic, viral and microRNA families. Nucleic Acids Res, 2021. 49(D1): p. D192–D200.

27. Burge, S.W., et al., Rfam 11.0: 10 years of RNA families. Nucleic Acids Res, 2013. 41(Database issue): p. D226–32.

28. Kwak, D., et al., Open-Phylo: a customizable crowd-computing platform for multiple sequence alignment. Genome Biol, 2013. 14(10): p. R116.

29. Pereira, R.d.L., et al., Increasing player engagement, retention and performance through the inclusion of educational content in a citizen science game, in The 16th International Conference on the Foundations of Digital Games (FDG) 2021. 2021, Association for Computing Machinery: Montreal, QC, Canada. p. Article 16.

30. Vance-Chalcraft, H.D., et al., Citizen Science in Postsecondary Education: Current Practices and Knowledge Gaps. Bioscience, 2022. 72(3): p. 276–288.

31. Edgar, R.C., MUSCLE: a multiple sequence alignment method with reduced time and space complexity. BMC Bioinformatics, 2004. 5: p. 113.

32. Minh, B.Q., et al., IQ-TREE 2: New Models and Efficient Methods for Phylogenetic Inference in the Genomic Era. Mol Biol Evol, 2020. 37(5): p. 1530–1534.

33. Waldispuhl, J., A. Kam, and P.P. Gardner, Crowdsourcing RNA structural alignments with an online computer game. Pac Symp Biocomput, 2015: p. 330–41.

34. Singh, A., et al., Lessons from an Online Massive Genomics Computer Game. Proceedings of the AAAI Conference on Human Computation and Crowdsourcing, 2017. 5(1): p. 177–186.

35. Fitch, W.M., Toward Defining the Course of Evolution: Minimum Change for a Specific Tree Topology. Systematic Biology, 1971. 20(4): p. 406–416.

36. Gaston, J. and S. Cooper, To Three or not to Three: Improving Human Computation Game Onboarding with a Three-Star System. Proc SIGCHI Conf Hum Factor Comput Syst, 2017. 2017: p. 5034–5039.

37. Blair, L., et al., Understanding the Role of Achievements in Game-Based Learning. International Journal of Serious Games, 2016. 3(4).

38. Fu, X. and J. Zhang, The influence of strategy video game and its background music on cognitive control. EC Psychology and Psychiatry, 2017. 2(1): p. 15–25.

39. Tremblay-Savard, O., A. Butyaev, and J. Waldispühl, Collaborative Solving in a Human Computing Game Using a Market, Skills and Challenges, in Proceedings of the 2016 Annual Symposium on Computer-Human Interaction in Play. 2016, Association for Computing Machinery: Austin, Texas, USA. p. 130–141.

40. Waldispuhl, J., et al., Leveling up citizen science. Nat Biotechnol, 2020. 38(10): p. 1124–1126.

41. Bonney, R., et al., Citizen Science: A Developing Tool for Expanding Science Knowledge and Scientific Literacy. BioScience, 2009. 59(11): p. 977–984.

42. Allf, B.C., et al., Citizen Science as an Ecosystem of Engagement: Implications for Learning and Broadening Participation. Bioscience, 2022. 72(7): p. 651–663.

