## Supplementary material for "When Online Citizen Science meets Teaching: Storyfication of a science discovery game to teach, learn, contribute to genomic research"

### When citizen science games meet teaching: a storified citizen science computer game to teach and contribute to genomic data analysis

Chrisostomos Drogaris, Alexander Butyaev, Elena Nazarova, Harsh Patel, Akash Singh, Breden Kadota, Jérôme Waldispühl

#### Architecture of the system

Figure 1 showcases the relationship between Phylo and OpenPhylo. Both Phylo and OpenPhylo use the same shared database. Phylo uses the same database of users to log-in as OpenPhylo. Phylo the game presents puzzle to users to solve. Once a user solves a puzzle, a solution record is created in the shared database.

Since the OpenPhylo server uses the same shared database, the OpenPhylo backend can utilize the solutions from the Phylo game. It can also create and edit existing puzzles. This shared database is a necessary and key component in providing the opportunity for OpenPhylo to provide useful services to supplement the Phylo game. The OpenPhylo server then communicates with the OpenPhylo frontend for users to interact with.

The OpenPhylo server is a Node.JS backend, while the OpenPhylo frontend is made with React.JS.

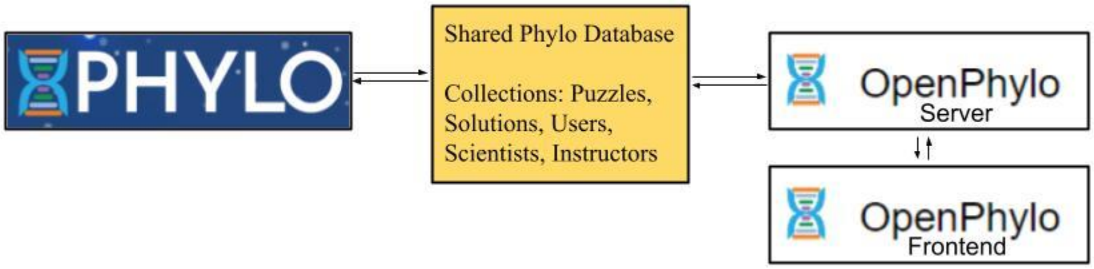


*Figure 1: Architecture of OpenPhylo.*

#### User Study

Between March 27th, 2022, and April 26th, 2022, we conducted a user study to evaluate the effectiveness of the new story mode as a teaching support activity. We asked 13 students (2 graduate students and 11 senior undergraduate students) registered in a graduate level bioinformatics course to complete the story mode of Phylo and monitored their progresses with the Open-Phylo instructor monitoring interface. Then, we ask them to complete an anonymous survey. All students completed the survey.

The survey included questions about the platform used (Web or mobile) as well a free text form that allowed the participants to leave us any comment they wanted. Eleven participants used the web version, and the two others uses the mobile version (iOS or Android). We do not report the free form answers. We also asked the participants to rate their experience using a 5 stars system. Two participants gave 5 stars, seven participants gave 5 stars, three participants gave 3 stars, and one participant gave 2 stars.

We present below the questions and distribution of answers that are discussed in the main article.

#### Questions about the story mode

- **What does the story bring to this citizen science game?**

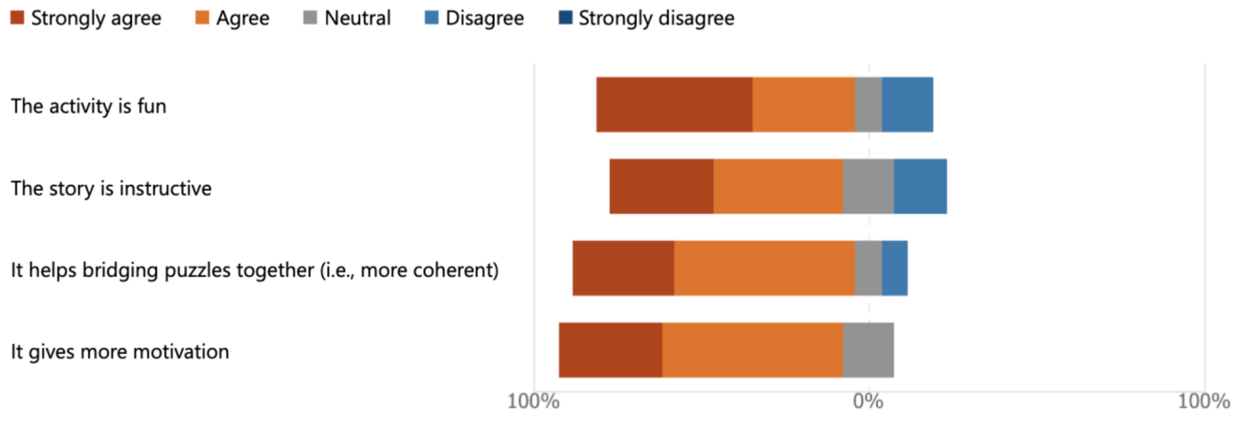


#### Structure and organization of the chapters

- **Please, indicate how much you agree with the following statements about Chapter 2 of the story (basic multiple sequence alignments)**

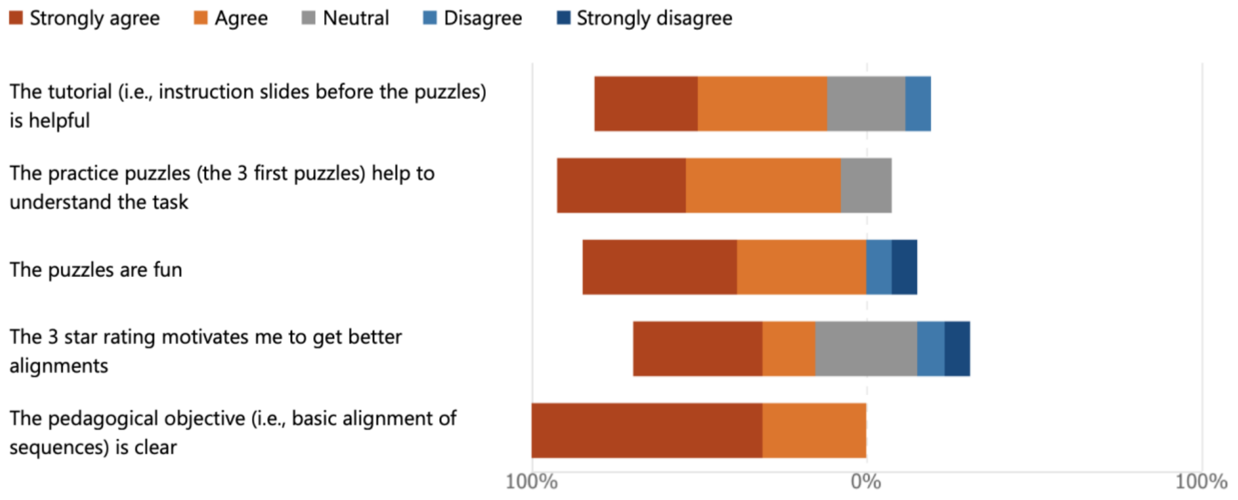

- **Please, indicate how much you agree with the following statements about Chapter 3 of the story (multiple sequence alignments with phylogeny)**

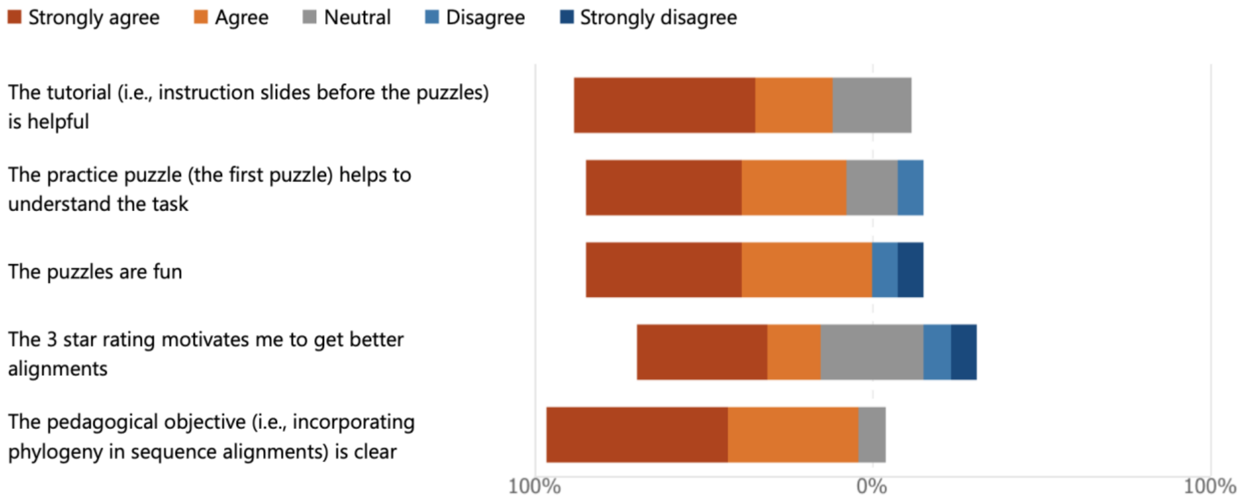


- **Please, indicate how much you agree with the following statements about Chapter 6 of the story (RNA alignments with secondary structures)**

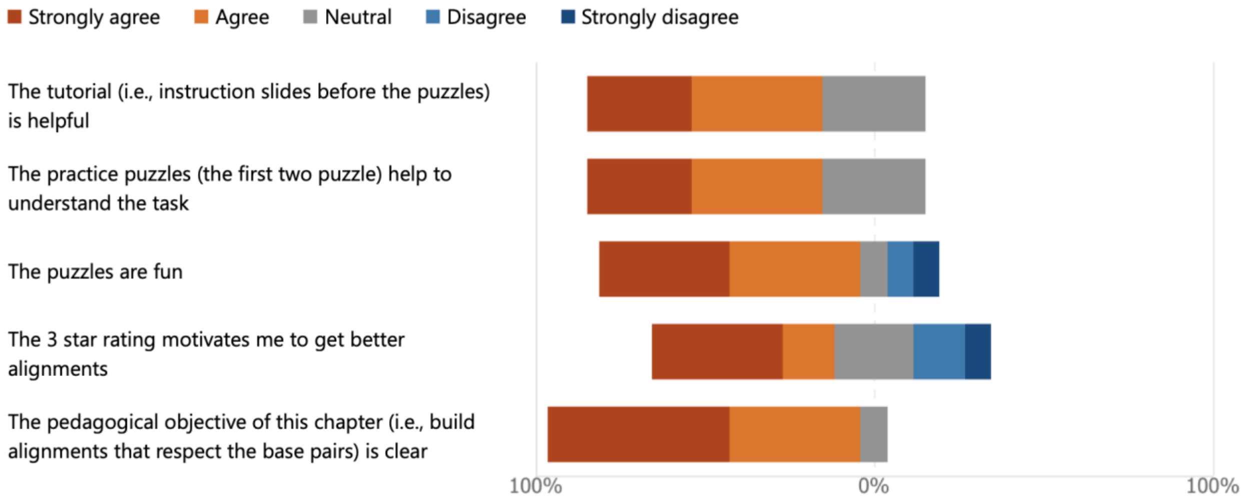

- **Please, indicate how much you agree with the following statements about Chapter 7 of the story (alignment of structured RNAs with base pair edits)**

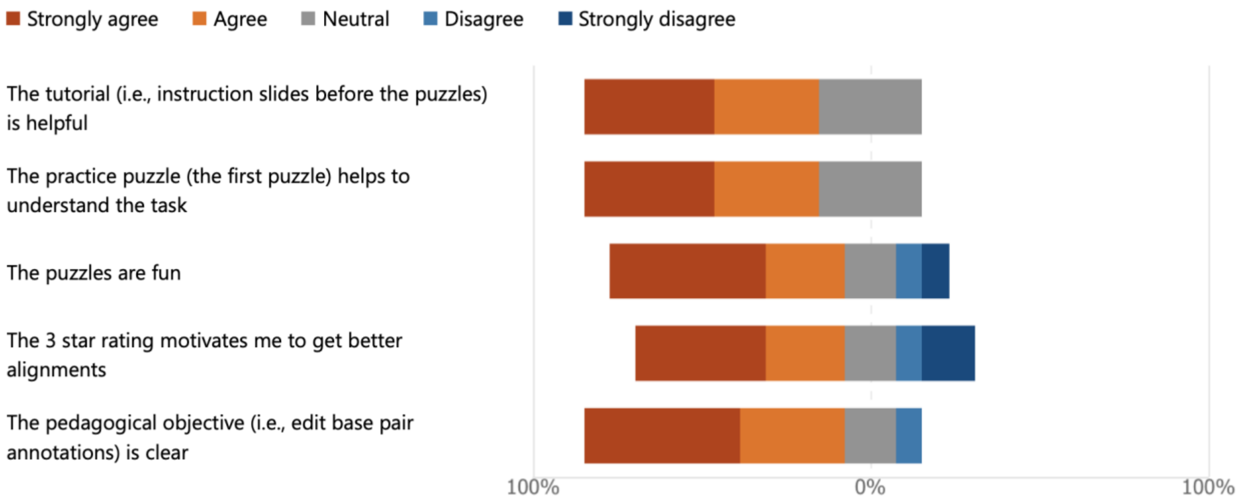
